# Brain State Dynamics and Developmental Differences in Reading Comprehension

**DOI:** 10.64898/2026.08.12.744344

**Authors:** Jia Zhang, Lanfang Liu, Jie Chen, Ningxin Zhao, Hehui Li, Xiujie Yang, Xiangzhi Meng, Guosheng Ding

**Affiliations:** School of Psychological and Cognitive Sciences, Beijing Language and Culture University, Beijing, 100083, China; Institute of Life and Health Sciences, Beijing Language and Culture University, Beijing, 100083, China; Key Laboratory of Language and Cognitive Science (Ministry of Education), Beijing Language and Culture University, Beijing, 100083, China; State Key Laboratory of Cognitive Neuroscience and Learning and IDG/McGovern Institute for Brain Research, Beijing Normal University, Beijing, 100875, China; Department of Psychology, School of Arts and Sciences, Beijing Normal University at Zhuhai, Zhuhai, 519085, China; Beijing Key Laboratory of Applied Experimental Psychology, National Demonstration Center for Experimental Psychology Education, Faculty of Psychology, Beijing Normal University, 100875, China; Center for Brain Disorders and Cognitive Sciences, School of Psychology, Shenzhen University, Shenzhen, 518060, China; School of Psychological and Cognitive Sciences and Beijing Key Laboratory of Behavior and Mental Health, Key Laboratory of Machine Perception (Ministry of Education), Peking University, Beijing, China, 100871; PekingU-PolyU Center for Child Development and Learning, Peking University, Beijing, 100871, China

**Keywords:** brain state dynamics, HMM, reading comprehension, reading development, fMRI

## Abstract

Reading comprehension is a complex cognitive task that involves dynamic interactions between the brain and external information. Previous studies on reading development primarily focused on localized or static brain activities. However, it remains an enigma how brain state dynamics evolve with development underlying reading comprehension. This study aims to address this issue by combining functional magnetic resonance imaging (fMRI) with Hidden Markov Model (HMM) to explore brain state dynamics. A total of 35 typically developing children and 31 adults were scanned while reading a story. Our results demonstrated a tripartite brain state organization, characterized respectively by high activities in the visual (State #1), language (State #2), and default mode network (DMN, State #3) regions. Children exhibited significantly longer dwell time in the DMN state (State #3) compared to adults, along with a higher probability of transitioning from the language state (State #2) to the DMN state (State #3). In addition, adults exhibited greater flexibility in state transitions during reading comprehension. Finally, the alignment between the dynamic states of children and the average states of adults was a significant positive predictor of their reading comprehension performance. This study provides a novel, intuitive perspective on how brain state dynamics evolve during the development of reading comprehension.

## 1. Introduction

Reading is a fundamental human capacity through which individuals acquire knowledge about the world and develop core cognitive abilities. Efficient reading comprehension is essential not only for academic achievement but also for social engagement and broader developmental outcomes (Hjetland et al., 2020). During reading comprehension, readers continuously extract and construct meaning from text by integrating explicit information with inferences derived from both textual cues and prior knowledge (Day & Park, 2005).

Despite its inherently dynamic nature, numerous studies on reading comprehension have focused on localized or static brain activities. Yet reading is a highly integrative process that relies on the flexible, time-varying coordination of multiple brain networks (Fedorenko & Thompson-Schill, 2014), which can be best captured by dynamic brain network approaches such as dynamic functional connectivity analysis (Wolff et al., 2022). To date, however, these methods have not been directly applied to reading comprehension, even though they successfully elucidated the neural dynamics of narrative understanding during spoken language processing and naturalistic paradigms such as movie watching (Liu et al., 2025; Song et al., 2021; Tang et al., 2023).

Dynamic functional connectivity goes beyond static analyses by characterizing how interactions among brain regions evolve over time. In particular, it enables the segmentation of brain activity into discrete, recurring states and reveals how the brain transitions between them (Hutchison et al., 2013). This framework is especially relevant to reading, which is not a static or passive process but one that requires continuous cognitive engagement, information integration, and flexible adaptation to textual complexities. Therefore, the present study prioritizes the investigation of brain state dynamics underlying reading comprehension.

Recent work by Liu et al. (2025) applied Hidden Markov Model (HMM; Rabiner & Juang, 1986) to speech comprehension in adults and identified a tripartite latent state space governing whole-brain dynamics. These states are characterized by elevated activity in (1) sensory-motor regions, (2) bilateral temporal language regions, and (3) the default mode network (DMN). This pattern aligns closely with the psycholinguistic model proposed by Berwick et al. (2013). According to this model, the core language faculty comprises syntactic rules and lexical representations, which interacts with two critical interfaces: an external sensory-motor interface (linking linguistic forms to the physical world) and an internal conceptual-intentional interface (connecting language to thought and intention).

Although reading and speech differed in input modality—reading relies on visual processing of orthographic symbols (e.g., via the visual word form area, VWFA), whereas speech depends on auditory parsing of acoustic signals, their core computations, such as semantic reasoning and integration, are largely shared (Perfetti & Stafura, 2014). This supports the nation of a modality-independent core language system (Berwick et al. 2013), suggesting that the reading comprehension should similarly engage the tripartite architecture involving sensory-motor, language, and DMN systems.

On the contrary, unlike speech comprehension, reading is a recent cultural invention for which the human brain did not evolve dedicated neural circuits. Instead, literacy recruit pre-existing cortical networks that are both functionally compatible and sufficiently plastic to accommodate new symbolic demands (Dehaene & Cohen, 2007). From the perspective of neuronal recycling, acquiring reading skills likely involve the functional reorganization of speech-related circuits to support visual word recognition (Dehaene et al., 2015; Hasson et al., 2016). Therefore, the brain state observed during reading comprehension may differ from the brain state involved in speech comprehension.

Moreover, reading development follows a well-established trajectory that transit from “learn to read” to “read to learn” through the formal education (Fox & Alexander, 2011). As reading becomes automatized, it becomes a powerful tool for ongoing knowledge acquisition and cognitive growth. Critically, reading experience shapes and modulates the brain’s reading network (Dehaene & Cohen, 2007). This modulation underpins the neural differences in reading between adults and children (Houdé et al., 2010; Martin et al., 2015; Zhu et al., 2014). Zhou et al. (2021) observed a developmental shift from phonological to semantic processing during reading, implying changes in the temporal dynamics of brain states during reading comprehension, particularly in dwell time and transitions among functional configurations.

Dynamic brain-state approaches, grounded in Dynamic Systems Theory (Smith & Thelen, 2003; Thelen & Smith, 1994), is ideally suited to capture such functional development. The brain is an intricate and dynamic system whose inter-regional interactions reorganized with maturation, resulting in more efficient and flexible information flow (Bassett & Sporns, 2017). Building on these insights, we hypothesize that: (1) Children will exhibit a higher transition probability from language-dominant states to the DMN, reflecting greater reliance on conceptual-intentional processing as they move beyond decoding toward meaning construction, and will show longer dwell time in the DMN. (2) Adults, with greater proficiency in contextual integration and complex reasoning, will demonstrate more flexible transitions across brain states, indicating enhanced network adaptability. (3) Greater alignment between a child’s brain state dynamics and the adult “ideal” pattern will predict better reading comprehension performance, suggesting that functional maturation underpins reading skill.

To test these hypotheses, we applied HMM (following Liu et al., 2025) to fMRI data collected during narrative reading in both children and adults. We quantified key dynamic metrics, including state transition probability, fractional occupancies, mean dwell time, and state switching rate. We further analyzed the topological properties of each brain state using graph theory methods. To determine whether the observed effect in whole-brain dynamics between children and adults is task-specific, we also compared whole-brain dynamics during resting-state scans between the two groups. By integrating dynamic network modeling with a developmental perspective, this study advances our understanding of the neural mechanisms underlying reading comprehension and offer a novel systems-level view of how brain flexibility and adaptability underpin literacy acquisition.

## 2. Methods and materials

### 2.1 Participants

A total of 35 typically developing children (mean age = 11.77 years, SD = 1.15, 22 males) and 31 typically developing adults (mean age = 23.58 years, SD = 2.41, 13 males) participated in the study. Children were fourth to sixth-grade students recruited from primary schools in Beijing. Adults were university students recruited from institutions in the same city. Eight children and two adults were excluded from the subsequent analysis due to excessive head movement (see details in 2.5). The final neuroimaging sample comprised 27 children and 29 adults.

All participants were native speakers of Mandarin Chinese, right-handed, and had normal or corrected-to-normal vision. None of them had a history of either neurological diseases or psychiatric disorders. Written informed consent was obtained from all adults and from both children and their legal guardians prior to testing. This study was approved by the Ethical Review Board of the State Key Laboratory of Cognitive Neuroscience and Learning of Beijing Normal University.

### 2.2 Behavioral measurements

#### Rapid automatized naming (RAN)

RAN was assessed to measure automatized access to phonological representation (Raberger and Wimmer, 2003), a process that shares cognitive resources with reading comprehension. When performing this task, the participants were instructed to read out visually-presented Arabic numbers as quickly and accurately as possible. The participants did the test twice, and the total naming time was recorded on each trial. The average of the two trials was calculated as the final score, with lower scores indicating better performance. Both children and adults completed this test.

#### Verbal fluency

Verbal fluency was evaluated using a semantic fluency paradigm, which assess the ability to retrieve lexical items under restricted search conditions (Lezak et al., 2004). Participants generated as many words as possible within one minute for each of four categories, including animals, fruits, work, and words containing the sound ‘fa’. The total number correct, non-repeated responses across four categories was averaged to yield fluency score. Higher scores reflected better verbal fluency. This task was completed by both children and adults.

#### Reading comprehension

Reading comprehension was evaluated using a standardized text comprehension test (Yu et al., 2022), with materials publicly available via the Dweipsy psychological research platform (www.dweipsy.com/lattice). The test consisted of sixty questions. For each question, children were asked to select the answer that best fits the context after reading a paragraph. The texts covered a variety of genres, including ancient poems, short stories, children’s songs, and novels. The number of correct answers was used as the raw score, with higher scores indicating stronger comprehension. Of the 27 children included in the final fMRI analyses, only 23 completed this test.

### 2.3 Experimental design

Participants performed a story comprehension task during fMRI scanning. The genre of the story was narrative, with a total duration of 4 minutes (240 seconds). There was a 45-second baseline before and after the story, during which the participants looked at a centered fixation point on the screen. The complete story consisted of 25 sentences, with each screen containing 18 ± 2 characters and being presented for 6 seconds. Before the experiment, the participants were instructed required to read the story carefully. After scanning, they were asked to retell the content of the story they had read in the scanner, and answer some reading comprehension questions. The questions included three types: detail recall (e.g., What musical instrument did Xiaoming play? A. Flute. B. Xiao. C. Sonar), inferential reasoning (e.g., Why did the boat capsize? A. The dolphins got too close and created waves. B. Due to weather, the sea was too rough. C. Other), and main idea identification (e.g., Provide an appropriate title for the story).

In addition to the task-based scan, resting-state fMRI data were collected, during which the participants were instructed to keep their eyes open, fixate on a central crosshair, remain still, and not focus on any specific thought.

### 2.4 Image acquisition

Brain images were obtained on a 3-T Siemens Prisma Scanner in the Imaging Center for Brain Research at Peking University. The parameters were as follows: flip angle (FA) = 65°; echo time (TE) = 30 ms; repetition time (TR) = 750 ms; field of view (FOV) = 200 mm × 200 mm; slice thickness = 2.5 mm, voxel size = 2.5 mm× 2.5 mm × 2.5mm, 55 slices, accelerate factor = 5, and interleaved slice acquisition. T1-weighted images were also acquired using the following parameters: FA = 7°; TE = 3.1 ms; TR = 2530 ms; FOV = 205 mm × 205 mm; slice thickness = 0.8 mm, voxel size = 0.8 mm× 0.8 mm × 0.8 mm, and 192 slices.

### 2.5 fMRI data acquisition and preprocessing

The preprocessing of functional magnetic resonance imaging (fMRI) data was conducted using SPM12 (https://www.fil.ion.ucl.ac.uk/spm/software/spm12/) and DPABI (https://rfmri.org/DPABI, Yan et al., 2016). First, a slice-timing correction was performed. Subsequently, spatial registration was conducted by first aligning each participant’s functional images to their structural images and then registering the structural images to the Montreal Neurological Institute (MNI) template, with a resolution of 2 mm × 2 mm × 2 mm. Then, linear trends removing and high-pass filtering (a cutoff of 1/128 Hz) were applied. Additionally, Friston-24 head motion parameters (Friston et al., 1996), white matter signal, and CSF signal were regressed out to reduce physiological noise. Among all participants, eight children and two adults were excluded due to excessive head movement (> 2 mm maximum translation or 2° rotation) or mean framewise displacement (FD) exceeding 0.2 mm (Power et al., 2012), leaving 27 children and 29 adults included for the final analyses.

### 2.6 Data analysis

#### 2.6.1 Whole-brain parcellation

The inference for brain dynamic states was conducted at the whole-brain network level using a nine-network atlas (Liu et al., 2025; Zhang et al., 2022). This nine-network parcellation was obtained by applying a state-of-the-art method proposed by (Ji et al., 2019), using data from the 64 participants engaged in speech comprehension (Liu et al., 2025). Because this nine-network atlas was generated from task-state data (rather than resting-state data) and was based on Chinese narratives, it is well-suited to our reading comprehension paradigm and participant population. A detailed schematic of the nine network partition is presented in the supplementary material (Fig. S1).

#### 2.6.2 Brain state inference using Hidden Markov Model

We applied a Hidden Markov model (HMM) to infer latent brain states based on the time courses of the nine networks during reading comprehension, using the HMM-MAR toolbox (https://github.com/OHBA-analysis/HMM-MAR). The HMM assumes that the observed neural dynamics arise from transitions among a finite set of latent states, each characterized by a distinct multivariate Gaussian distribution.

For each participant, we first computed the functional connectivity matrices across nine networks. Prior to infer latent brain states, the network time courses were standardized within each participant. Then, we concatenated the standardized time courses across all 56 participants (27 children and 29 adults). of both the populations of children and adults were, yielding a data matrix of dimensions 56 subjects × 200 time points × 9 networks. This concatenation enabled group-level HMM estimation, ensuring consistent state definitions.

To determine optimal number of latent states (K), we evaluate models with K ranging from 2 to 10. Model selection was guided by two clusters validity indices: the Calinski-Harabasz (Caliński & Harabasz, 1974; Liu et al., 2025) and the silhouette (Rousseeuw, 1987; Tang et al., 2023). Both metrics were widely used in combination to identify the optimal value of K (Ge et al., 2024; Liang et al., 2020; Shi et al., 2023; Xing et al., 2022; You & Park, 2022), with higher scores indicating better clustering performance. For each participant, we computed both indices and then averaged them across all participants. To combine these two criteria, we converted each into Z scores and then summed them to form a composite score. The K yielding the highest composite score was selected as optimal.

Using this optimal K, we estimated the HMM and extracted, for each participant, the posterior probability time course for each state. From these, we derived three key dynamic metrics per participant (Vidaurre et al., 2017): fractional occupancies (FO), mean dwell time (MDT), and switching rate (SR). Specifically, fractional occupancies of each state are computed as the ratio of the activated HMM states across the time courses. The dwell time of each state is the duration of the visits to that particular state. The switching rate are defined as the frequency of transitions between different states. Group differences in these metrics between children and adults were assessed using two-tail independent-sample *t*-tests, with multiple-comparison correction (False Discovery Rate-FDR, α = 0.05). A schematic overview of the analysis pipeline is shown in Fig. 1.

**Fig. 1.**
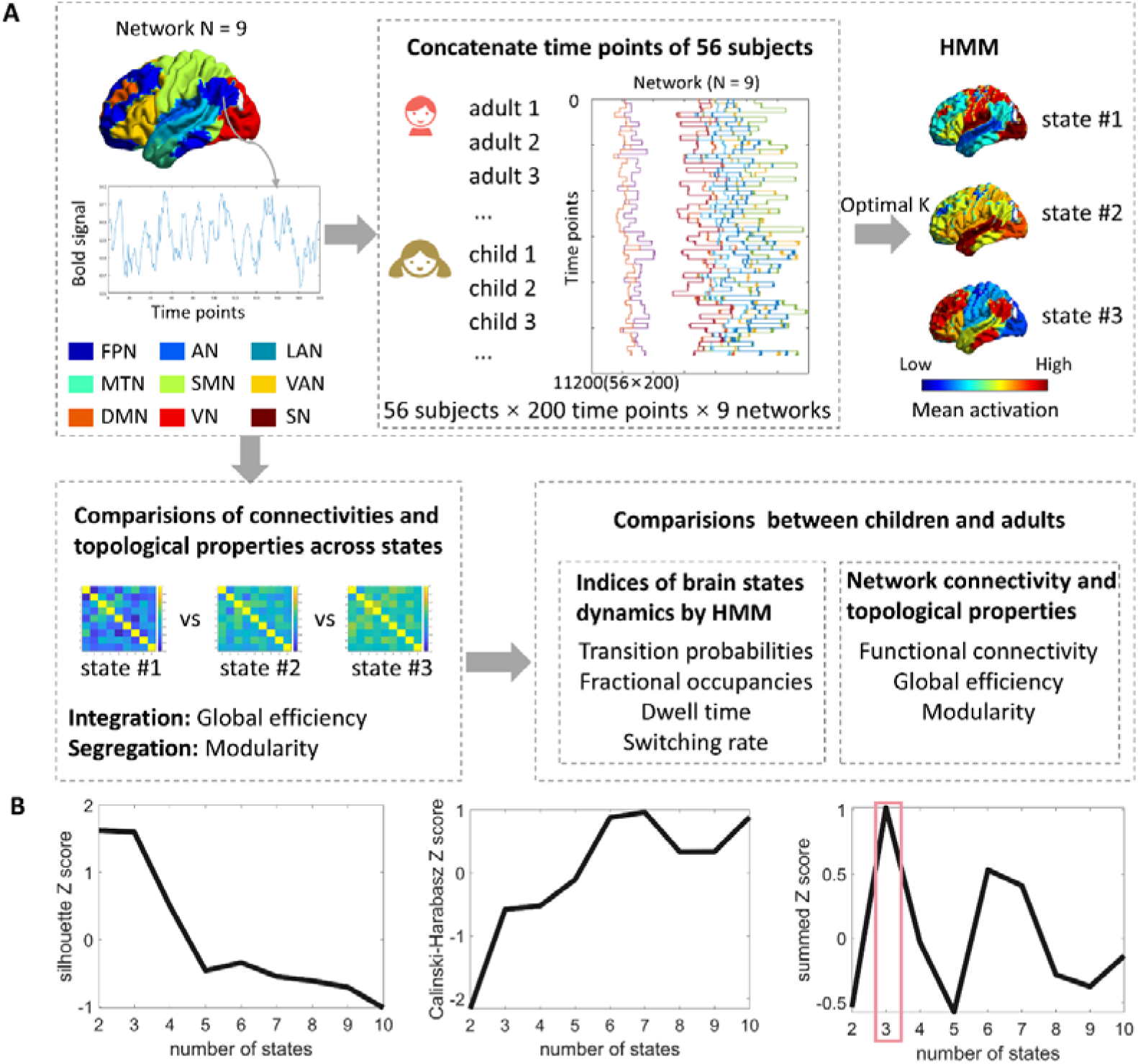
(A) Schematic illustration of the analyses of brain dynamic states. (B) Identify the optimal number of brain states based on two clustering performance indices (as quantified by the summed Z scores).

As a control analysis, we applied the same HMM with identical K value to resting-state data to determine whether the observed developmental differences were specific to the reading task or merely reflected task-independent maturational effect. If group differences are still observed during the resting state, it suggests that the developmental differences were intrinsic, not specific to the reading task. Conversely, if these differences disappear during the resting state, they would support task-selectivity.

#### 2.6.3 Analyses of network connectivity and topological properties

We firstly obtained the functional connectivity of each subject under three brain states and conducted the test on the differences across three states. Subsequently, we compared the differences in the functional connectivity between networks of adults and children under each state. Next, we calculated the functional integration and segregation of the whole brain under each state using the graph theoretical analysis. We constructed a weighted and undirected graph for each participant under each state. Specifically, the nine networks were regarded as nodes, and the connections between time courses corresponding to specific states were regarded as edges, which were operated by Brain Connectivity Toolbox (Whitfield-Gabrieli & Nieto-Castanon, 2012). Functional segregation is characterized by weakened interregional connections between networks and was measured by the network modularity of brain networks using Louvain algorithm with a resolution parameter gamma = 1 (Liu et al., 2025). Functional integration refers to the capacity of interactions between segregated regions and was quantified by the global efficiency of brain networks (Rubinov & Sporns, 2010). Then, *t*-tests were employed to examine the differences of the global efficiency and modularity across three states. For each state, we also performed two-tail two-sample *t*-tests for the global efficiency and modularity between children and adults (False Discovery Rate-FDR, α = 0.05). For each state, beyond computing a subject-level functional connectivity (FC) matrix, the HMM model generated a group-level FC matrix. For validation, we additionally computed these indices using the group-level FC matrices (each state corresponds to one FC matrix) estimated by the HMM.

#### 2.6.4 Correlation of latent state dynamics with reading comprehension

To assess the functional relevance of the brain state dynamics to reading skill we examined whether greater similarity in state transition patterns between children and adults was associated with better reading comprehension performance. We hypothesized that adult-like brain dynamics, which could be defined as alignment with an “ideal” adult state trajectory, supports efficient comprehension. To test this hypothesis, we quantified the alignment of brain state fluctuations between each child and each adult, then averaging these values (each child to all adults) to obtain an inter-subject correlation (ISC) index. This index was then used to predict children’s reading comprehension scores. To control for potential confounds related to motion, we computed ISC in head movement trajectories, quantified by FD during fMRI scanning, and included these as covariates in a partial correlation analysis.

## 3. Results

### 3.1 Demographic and behavioral performance

There were no significant differences in gender (χ^2^ = 2.89, *p* = 0.09) between children and adults. Compared with children, adults had significantly better rapid automatized naming (*t* _(64)_ = 4.65, *p* < 0.001) and verbal fluency (*t* _(64)_ = - 4.86, *p* < 0.001) scores (Table 1).

**Table 1.**
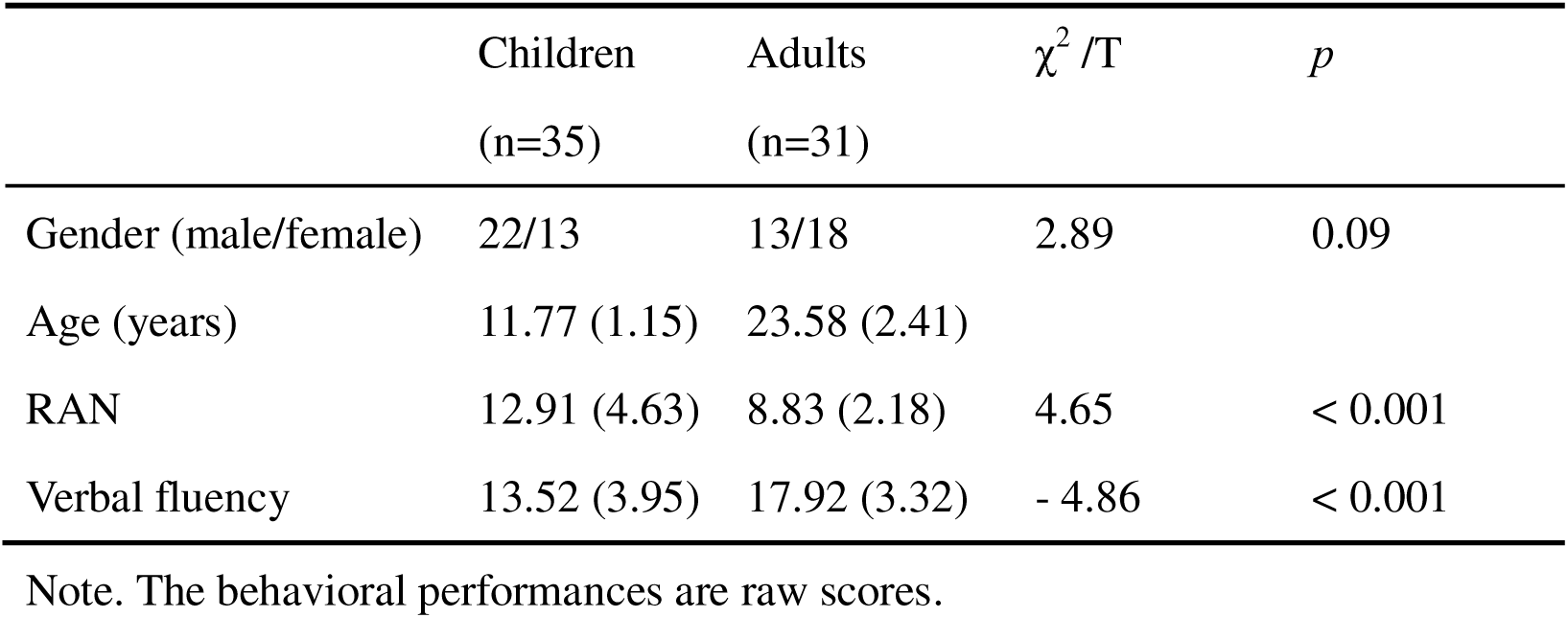
Demographics and behavioral performance.

| | Children<br>(n=35) | Adults<br>(n=31) | $\chi^2$ /T | *p* |
| --- | --- | --- | --- | --- |
| Gender (male/female) | 22/13 | 13/18 | 2.89 | 0.09 |
| Age (years) | 11.77 (1.15) | 23.58 (2.41) |  |  |
| RAN | 12.91 (4.63) | 8.83 (2.18) | 4.65 | < 0.001 |
| Verbal fluency | 13.52 (3.95) | 17.92 (3.32) | - 4.86 | < 0.001 |

Note. The behavioral performances are raw scores.

### 3.2 Tripartite brain state organization during reading

To infer the latent brain states, we concatenated the time courses of the BOLD signals obtained from both children and adults. Across a range of candidate models with K values from 2 to 10, the silhouette generally decreased as K increased, while the Calinski-Harabasz showed an increasing trend. The HMM model with K = 3 demonstrated the best overall performance, as quantified by the summed Z scores (Fig. 1B). The HMM model generated, for each state, a group-level activation map across nine brain networks and a FC matrix representing the interactions between these networks. We identified tripartite brain states, each exhibiting unique activity patterns corresponding to the three language components, consistent with the theoretical framework proposed by Berwick et al. (2013). We further computed the similarity of activity patterns between the three states in our study and those reported by Liu et al. (2025), as detailed in Fig. S2.

We found that the first state (State #1) was characterized by high activities in the visual, auditory and somatomotor networks, corresponding to the external sensory-motor module. The second state (State #2) was dominated by the language and medial-temporal networks, reflecting core linguistic processing. The third state (State #3) showed strong activation of the DMN and frontal-parietal networks, consistent with the internal conceptual-intentional component (Fig. 2). Our results demonstrated the applicability of the tripartite latent state space in reading comprehension.

**Fig. 2.**
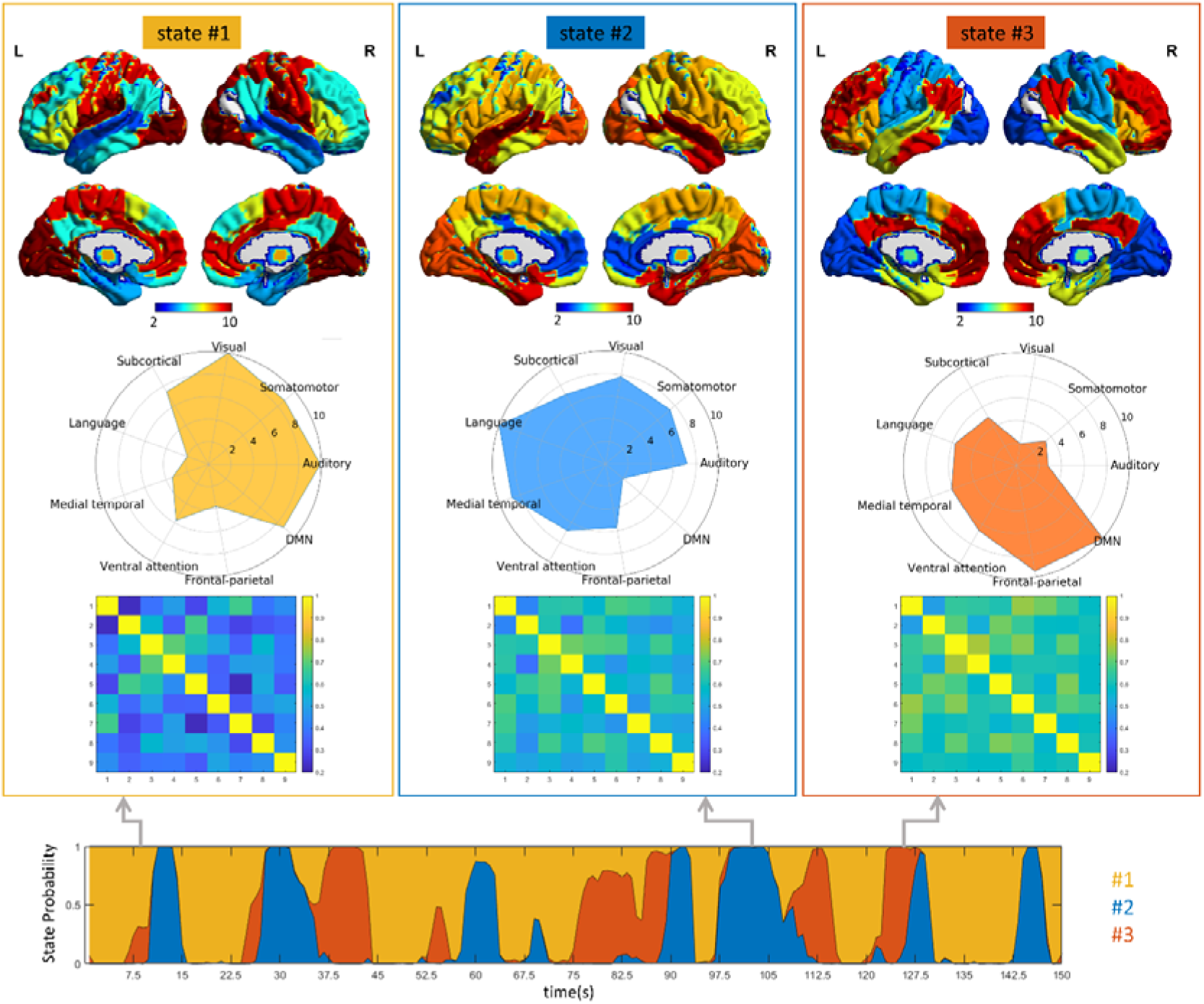
The group-level activation map on the nine networks and the functional connectivity matrix between these networks that HMM model generated for each state. For visualization purpose, the spatial map was normalized to the range [2,10] with min-max normalization. The numbers from 1 to 9 represent the frontal-parietal network, the auditory network, the language network, the medial temporal network, the somatomotor network, the ventral attention network, the default mode network, the visual network, and the subcortical network, respectively.

For the functional connectivity matrix HMM model generated, an averaged FC across nine networks within each state were then computed. The results showed that State #3 exhibited the highest FC (mean = 0.631, SD = 0.066), followed by State #2 (mean = 0.568, SD = 0.083), while State #1 had the lowest FC (mean = 0.432, SD = 0.121). These results suggest that brain communication is relatively weak in the sensory-motor module compared to the other two states.

### 3.3 Brain network connectivity and topological properties: state differences and child-adult comparisons

We first compared the values of the inter-network connections of all subjects across three states. The results revealed that inter-network connectivity in State #3 was greater than that in the other two states, and inter-network connectivity in State #2 also exceeded that observed in State #1 (FDR *ps* < 0.05). Additionally, the connections between auditory network and frontal-parietal, ventral attention network, the connections between ventral attention network and somatomotor, visual network were observed in all pairs of comparison, demonstrating gradually increased from State #1 to State #3, suggesting the changes in the functional connectivity between the low-level somatomotor network and the high-level association cortex across different states. (Fig. 3A). Subsequently, for each state, we conducted two-sample *t*-tests on the inter-network connections between adults and children. The results showed that differences between children and adults were only found in State #2. Specifically, the connections between the DMN and the ventral attention, and between DMN and subcortical regions were higher in children than in adults (Fig. 3B). Such effects between adults and children were also obtained using static networks (Fig. 3C).

**Fig. 3.**
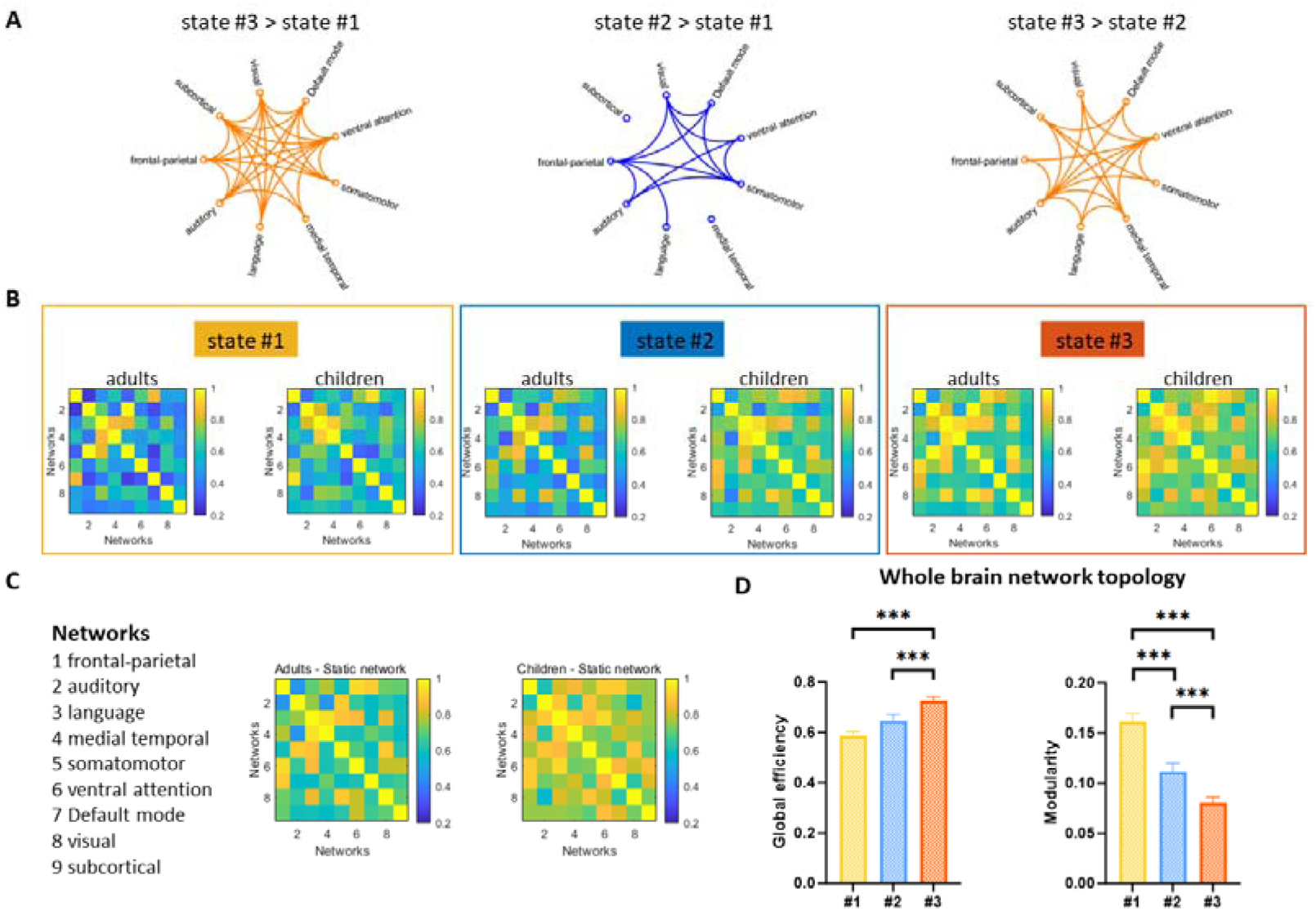
Network connectivity and topological properties among three brain states. (A) Network connectivity across three brain states in all subjects. (B) The functional connectivity patterns between networks for children and adults under different states using state-specific time courses from individual participants. (C) The static functional connectivity patterns for children and adults between different networks. (D) Topological properties of whole-brain networks across three brain states. State #3 demonstrated the highest global efficiency and the lowest modularity. The numbers from 1 to 9 represent the frontal-parietal network, the auditory network, the language network, the medial temporal network, the somatomotor network, the ventral attention network, the default mode network, the visual network, and the subcortical network, respectively.

We then compared which state demonstrated the highest degree of information integration. Based on the Hidden Markov Model (HMM), we obtained the time points at which each subject belongs to States #1, #2, and #3 respectively. We then extracted the time points that belong to each specific state, and thus, a functional connectivity matrix was obtained for each subject in each state. The graph theoretical analyses were subsequently utilized to assess the functional integration (i.e., global efficiency) and functional segregation (i.e., modularity) of the whole-brain networks. Specifically, state #3 showed the significantly higher global efficiency than state #1 and state #2 (*t* values > 3.54, *ps* < 0.001), while showed significantly lower modularity than that in state #1 and state #2 (*t* values > 3.9, *ps* < 0.001). These results suggest that the information integration is the highest in state #3 (Fig. 3D). In addition, the HMM model generated a functional connectivity matrix for each state (Fig. 2). We also calculated two graph theoretical indices (i.e., global efficiency and modularity) using the state-specific functional connectivity matrices estimated by the HMM. The global efficiency was highest on State #3 (0.631), next on State #2 (0.568) and the lowest on State #1 (0.432). An opposite pattern was found in network modularity. The modularity was highest on State #1 (0.142), next on State #2 (0.077) and the lowest on State #3 (0.061). These results showed that either using state-specific time courses from individual participants to construct the FC matrix or taking the FC matrix derived from the HMM were consistent. We also concerned whether the differences of network integration and segregation exist between children and adults using state-specific time courses from individual participants. However, we did not observe any significant differences between children and adults.

### 3.4 Comparisons of brain state dynamics between children and adults

We calculated the state transition probabilities, fractional occupancies, mean dwell time, and switching rate for each participant. In children, the brain spent most of the time on State #3 (mean FO = 42.1%), next on State #2 (mean FO =30.9%) and the least on State #1 (mean FO = 27.0 %). The same pattern was found in the dwelling time, with a group mean of 7.60s for State #3, 6.07s for State #2, and 5.82s for State #1. In adults, the brain spent more time on State #3 (mean FO = 34.9%) and State #2 (mean FO =34.3%) and the least on State #1 (mean FO = 30.8 %). As the dwelling time, the most was on State #2 (5.80s), next on State #3 (5.60s) and the least on State #1 (5.44s).

A 2×3 ANOVA was conducted with the state (#1 /#2 /#3) as the within-group factor and group (children/adults) as the between-group factor. For fractional occupancies, we found the main effect of state (*F* _(2,108)_ = 4.127, *p* = 0.019, *η*_p_^2^ = 0.071), with significant differences between state #3 and state #2. For dwell time, no significant effect was observed.

Then, we performed two-sample *t*-tests between children and adults. No significant result was found between children and adults for fractional occupancies. The mean dwell time in State #3 was greater in children than adults (*p* = 0.024, FDR corrected). Additionally, adults exhibited higher switching rate between three states (*p* = 0.002), which may indicate greater flexibility in state transitions during reading comprehension (Fig. 4B). Finally, for the transition probabilities, we found that children were more likely to transition from State #2 to State #3 compared to adults (*p* = 0.013, FDR corrected). In contrast, adults were more likely to transition from State #3 to State #2, which reached marginal significance (*p* = 0.056, FDR corrected) (Fig. 4A).

**Fig. 4.**
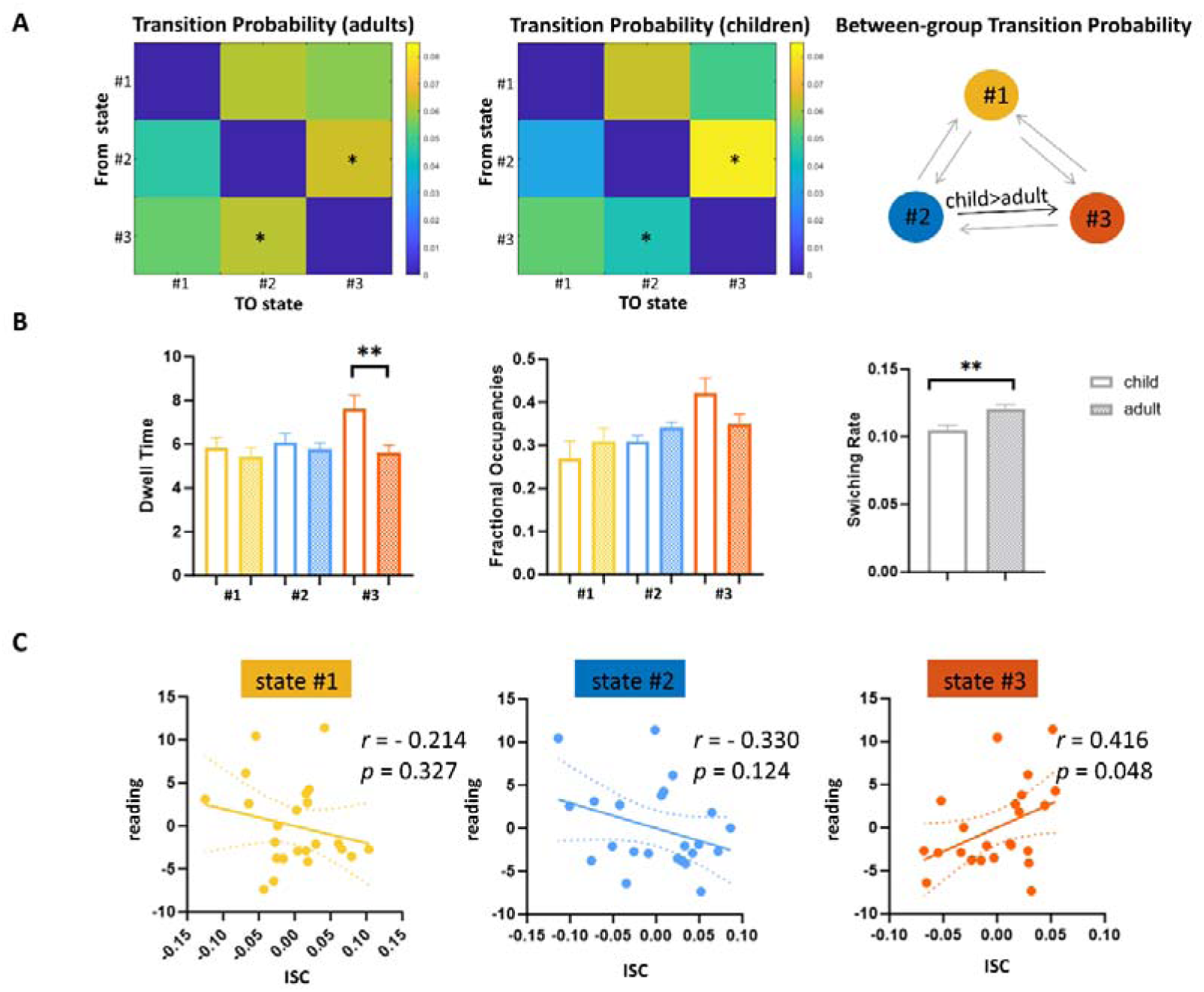
Differences of brain states dynamics between children and adults. (A) The differences of state transition probabilities. (B) The differences of the fractional occupancies, the mean dwell time, and the switching rate. (C) The alignment between each child and the average dynamic states of adults correlated with reading comprehension score. Y-axis represents the residual of reading comprehension score; X-axis represents the residual of the alignment with adults.

We also test whether individual differences in the fractional occupancies and the mean dwell time of latent states, and switching rate were associated with individual difference in reading-related skills (i.e., RAN, verbal fluency and reading comprehension) for children and adults, respectively. No significant result was found on any of the three states (*r* values < 0.30, *ps* > 0.11). Taken together, these findings suggested that the differences between adults and children observed in the overall magnitude of engagement in those states may be unrelated to the differences in reading-related skills.

### 3.5 Correlation of latent state dynamics with reading comprehension

We hypothesized that temporal alignment between a child’s brain state fluctuations and the adult “ideal” pattern would predict comprehension ability. To test this hypothesis, we calculated the alignment for each child by computing the Pearson correlation between their state expression probability time courses and that of each adult, followed by averaging these correlations. Subsequently, a Pearson correlation was computed between the inter-subject correlation (ISC) scores and children’s comprehension scores.

The results revealed that average alignment between each child’s state expression probability time courses and that of adults significantly positively predicted children’s reading comprehension performance, even with ISC in the FD during the fMRI scanning as the nuisance covariates. Notably, this effect was specific to State #3 (*r* = 0.416, *p* = 0.048, Fig. 4C), suggesting that children who transition into specific brain states (DMN) with timing similar to adults exhibit better reading comprehension abilities. These findings underscore the importance of the brain’s ability to promptly switch to specific states as a critical factor for effective reading comprehension.

### 3.6 No difference between children and adults of brain state dynamics during rest

To determine whether observed child–adult differences reflect task-specific or intrinsic developmental effects, we applied the same HMM (K = 3) to resting-state data from the same participants. The results revealed three states with moderate similarity in activity patterns to that of the reading comprehension condition (overall *r* _(25)_ = 0.489, *p* = 0.014, Fig. S3). We did not observe any significant differences between children and adults in transition probabilities, fractional occupancies, mean dwell time, and switching rate (FDR corrected, Fig. 5B). In addition, the average alignment between each child’s state expression probability time courses during rest and that of adults cannot predict children’s reading comprehension performance (*ps* > 0.219, Fig. 5C). Totally, our results suggested that the differences of tripartite latent space of whole-brain dynamics between children and adults were mainly driven by the reading task, rather than reflecting general maturational differences in intrinsic brain organization.

**Fig. 5.**
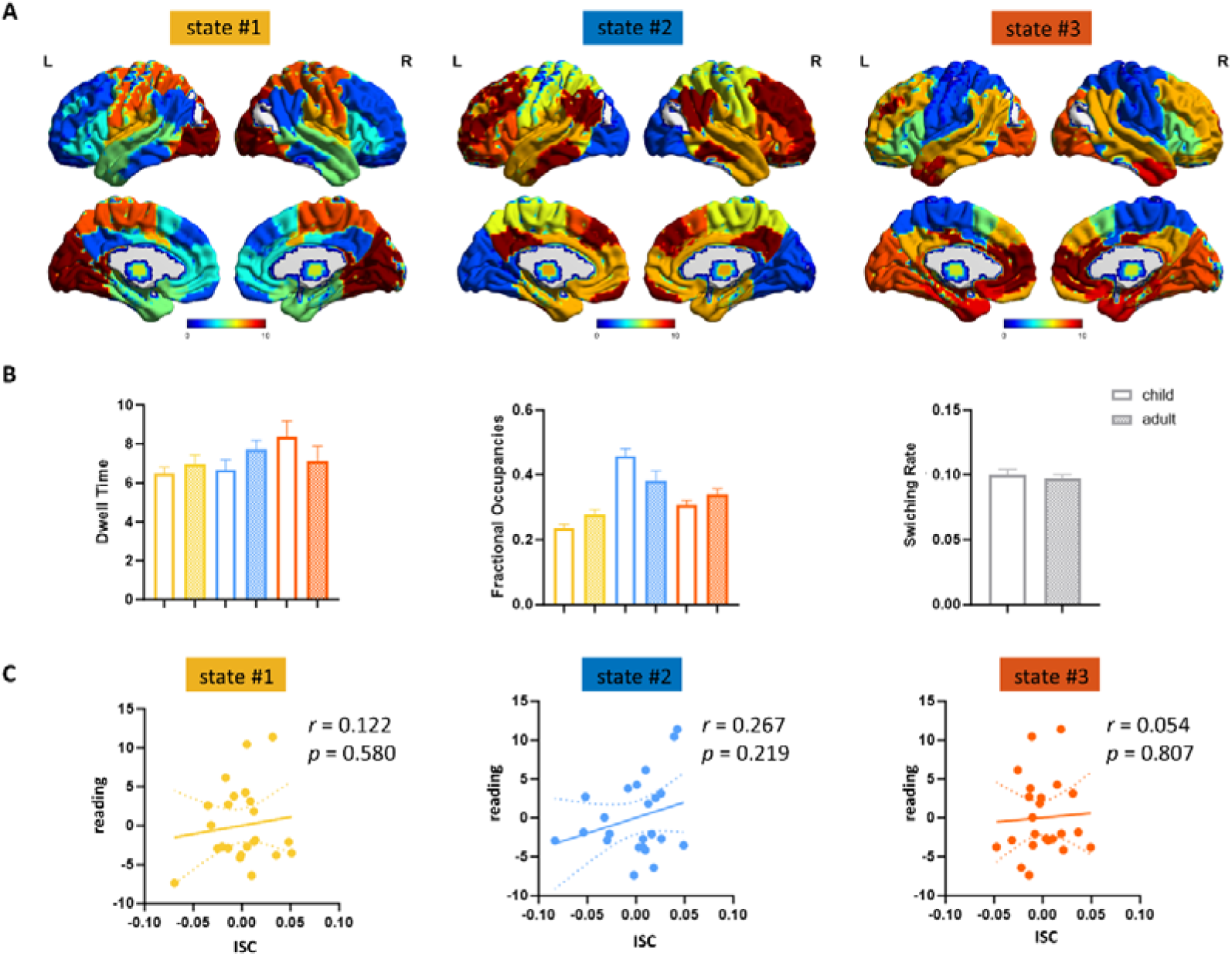
Brain state dynamics during rest. (A) During rest, the activity patterns of latent states were moderate similar to those during reading comprehension. (B) No differences of the mean dwell time, the fractional occupancies, and the switching rate were observed between children and adults. (C) The alignment between each child during rest and the average dynamic states of adults correlated with reading comprehension score. Y-axis represents the residual of reading comprehension score; X-axis represents the residual of the alignment with adults.

## 4. Discussion

Reading comprehension is a sophisticated cognitive task that requires an interactive dynamic between the brain and external information. Our study explored the brain state organization and its development in reading comprehension. Applying HMM, we found that the brain showed tripartite state space similar to that of speech comprehension, which were characterized respectively by high activities in the visual (State #1), language (State #2), and DMN (State #3) regions. We also observed functional connectivity and functional integration gradually increased from sensory-motor module (State #1) to default mode network (State #3), either using state-specific time courses from individual participants to construct the FC matrix or taking the FC matrix derived from the HMM. Compared to adults, the mean dwell time in DMN (State #3) was greater in children. And children are more likely to transition from the language state (State #2) to the DMN state (State #3). Adults demonstrated greater flexibility in state transitions during reading comprehension. Furthermore, the alignment between each child’s dynamic states and the average dynamic states of adults within DMN state significantly predicts their reading comprehension performance. Finally, by comparing the reading comprehension task with resting conditions, we found that the differences between children and adults were primarily observed during the reading task. This study provides insight into a more intuitive depiction of how the brain transitions between different states during development of reading comprehension.

Language is a complex cognitive process that requires consideration of the brain’s functionality as a whole. Hagoort (2014) also argued that the function of a brain region should not be understood from a localizationist perspective, but rather through the interactions of different networks. Therefore, a larger-scale connectivity perspective might be a more informative method for studying reading development. Graph theory, a method used extensively of late, makes it possible to understand functional connectivity in a large-scale network. In this approach, the human brain is proposed to be structurally and functionally organized into a complex network to facilitate the effective segregation and integration of information processing (Sporns, 2013). This network is described as a graph with nodes (brain regions) and edges (functional or structural connections, Bullmore & Sporns, 2009). Graph theory provides a powerful statistical framework for characterizing the development of brain systems in a comprehensive manner, considering not only relationships within a given system, but also how these relationships are situated within wider network contexts (Power et al., 2010). However, static network primarily captures the overall connectivity features of the brain during a specific task. For story reading, which have the temporal dimension, different brain functional network states may be activated. Therefore, compared to static graph theory analysis, dynamic network analysis divides brain functional networks into multiple states to capture the brain’s state at different time points, demonstrating the transitions between different brain states more intuitively (Hutchison et al., 2013). It’s more suitable for researchers to examine the connectivity patterns between brain regions at various time points and how these patterns change as the narrative unfolds during reading comprehension.

Recently, Liu et al. (2025) used HMM to reveal that, during speech comprehension, whole-brain networks predominantly oscillate within a tripartite latent state space, characterized by high activity in the sensory-motor (State #1), bilateral temporal (State #2), and DMN (State #3) regions, approving three modules in a psycholinguistic model (Berwick et al., 2013). Reading comprehension also leads to the widespread co-activation of brain regions, which involve not only the decoding of words, but also higher-order cognitive functions such as semantic reasoning and integration (Perfetti & Stafura, 2014). According to the model proposed by Berwick et al. (2013), the language system consists of three core components: syntactic rules and representations, the external sensory-motor interface, and the internal conceptual-intentional interface. Readers must continuously interact with information from the external world, along with extracting and constructing meaning from the text, using textual cues and knowledge to extract explicit information and infer implicit meaning. Effective reading comprehension requires continuous cognitive engagement, involving dynamic information processing and integration between external stimuli and internal neural representations. (Day & Park, 2005). Theoretically, the reading comprehension should also conform to the model proposed by Berwick et al. (2013). Our study supported that tripartite organization can serve as a principal applicable to language comprehension, whether in the auditory or visual modality.

Language development is a complex and dynamic process. Larsen-Freeman (1997) proposed that language learning is not a linear process solely driven by input but rather a dynamic system characterized by regression, stagnation, and even leaps forward. Several studies have demonstrated the utility of dynamic functional connectivity (FC) methods in understanding functional neurodevelopment. A study on participants aged 9 to 32 found that, while the brain’s functional connectivity topology remained stable across different age groups, the frequency and dwell time of certain functional connectivity states exhibited age-related variations during cognitive control tasks. Additionally, older individuals showed a higher frequency of state transitions (Hutchison & Morton, 2015). Other studies have also indicated that the connectivity between brain networks becomes increasingly variable with age (Marusak et al., 2017; Qin et al., 2015), leading to the hypothesis that improvements in cognitive function are supported by enhanced neural temporal dynamics (Hutchison & Morton, 2016). To date, there has been no studies investigating how brain states evolve with reading experience during reading comprehension. Therefore, it is crucial to examine the differences in brain-state dynamics between adults and children during reading comprehension. This study investigates the fractional occupancies, the mean dwell time, and switching rate across different states in adults and children. Our findings, using this dynamic brain state approach, showed that children exhibited a higher transition probability from the language state to the DMN state than adults, and with a longer dwell time in the DMN state. This may suggest that children are more reliant on transition from linguistic processing to information understanding during reading comprehension. Furthermore, during the development of reading comprehension, adults are able to more quickly establish connections between the storyline and semantics, compared to the reliance on basic phonological processing modules observed in children. This shift plays a crucial role in the development of reading skills (Kintsch & van Dijk, 1978). Consequently, we observed that adults exhibited greater switching rates, which may reflect their greater flexibility and more efficient information integration during reading comprehension, supporting the hypothesis that cognitive function improvement is driven by enhanced neural temporal dynamics. Based on Dynamic Systems Theory, our study provides a significant perspective for understanding brain development, the maturation of neural networks, and children’s performance on cognitive tasks.

One core concept in evolutionary psychology is that species gradually develop the “optimal” physiological and cognitive mechanisms through evolution to adapt to their environment. The structure and function of the brain have been continuously optimized throughout evolution to meet the cognitive demands of complex environments (Dunbar, 1998). From this perspective, the adult brain activity pattern can be considered an “ideal state”, as it represents the most efficient mode of the brain when confronted with complex tasks such as language comprehension and decision-making. Over the course of human evolution, the cognitive mechanisms and functional networks of the adult brain have undergone long periods of natural selection, enabling adults to perform cognitive tasks with greater efficiency and flexibility. Therefore, the gradual maturation of children’s brain and its progression towards the adult brain state can be understood as a process of development towards this “ideal state”. From this standpoint, the greater the alignment of a child’s brain activity with that of adults, the better their cognitive performance should be. Interestingly, our study found the more children are adults alike in alignment within the DMN state, the greater their reading comprehension performance would be. This result suggested that brain’s ability to promptly switch to specific states could be a critical predictor for reading comprehension ability. In line with our findings, Zhou et al. (2021) also found that the more the functional connectivity pattern of adults resembles that of children, the poorer their reading performance becomes. In other words, the gradual alignment of a child’s neural activity pattern with the adult level could indeed be a hallmark of cognitive development. This process likely reflects the maturation of cognitive functions following both developmental and environmental influences on the brain.

Interestingly, we found that the differences between children and adults were primarily concentrated in the DMN state. The maturation of the DMN may be a key factor driving these differences, as the functional connectivity of the DMN in children may not yet be fully developed, leading to their longer dwell time in this state (Fan et al., 2021). This prolonged dwell time in children may reflect their process of establishing and optimizing reading comprehension abilities, representing a core mechanism underlying the development of reading skills. The higher transition probability between the language state and the DMN state in children may indicate that they rely more on the DMN for semantic integration during reading comprehension, which is a high-level cognitive processes. This suggests that their brain networks have not yet achieved the efficiency and flexibility observed in adults. The differences in dynamic state indices between the two groups suggest that children and adults may adopt different cognitive strategies during reading comprehension. Children are more likely to frequently connect story information with their personal experiences, requiring greater activation of the DMN to support narrative simulation and semantic reasoning. In contrast, adults may rely on more streamlined strategies, integrating information efficiently through flexible state transitions. Furthermore, the correlation of latent state dynamics with behavioral performance revealed that the dynamic properties of the DMN might serve as a critical indicator of reading ability. The greater flexibility exhibited by adults likely reflects their established proficiency in reading skills, whereas the alignment of children’s dynamic states with the average dynamic states of adults may indicate that they are in the process of learning and emulating efficient neural network organization.

There are several limitations in our study. First, this research is an exploratory investigation into the development of brain states during reading comprehension. The reading comprehension task involved a relatively short story, with only 200 time points. Future studies could use longer time points for validation and examine different types of texts. Additionally, our study identified tripartite brain states highly similar to those observed by Liu et al. (2025) in speech comprehension, suggesting that the tripartite states may serve as a fundamental principle for language comprehension, regardless of whether the modality is visual or auditory. However, our study included only one story, which limits the depth of exploration on this issue. Future research could systematically examine whether the division of language comprehension into three stable states is independent of the modality of stimulus presentation.

## 5. Conclusion

Using a dynamic systems approach based on Hidden Markov Modeling, we reveal that reading comprehension is supported by a tripartite latent state space, consistent with the model proposed by Berwick et al. (2013). Developmental differences between children and adults were most pronounced in the DMN-dominated state, with children showing longer dwell time and stronger transitions into this state. Crucially, the alignment between a child’s brain state dynamics and the adult “ideal” pattern is a significant predictor of children’s reading comprehension performance. Moreover, these developmental differences were specific to the reading task and were not observed during rest. Together, these findings illustrate that reading development involves not just physiological maturation, but the refinement of dynamic network reconfiguration. The incorporation of time-resolved brain state modeling offers a powerful framework for understanding how the brain achieves flexible, efficient comprehension, and how this capacity emerges through development.

## Acknowledgements

This work was supported by grants from the National Natural Science Foundation of China (NSFC: 32371141, 32471100, 32500941). This work was also supported by the Youth Foundation for Humanities and Social Sciences Fund of Ministry of Education of China (grant number 25YJC740070).

## Ethics Approval Statement

We confirmed that the appropriate ethical approval has been received.

## Supplementary Materials

Supplementary material associated with this article can be found, in the online version.

