## Supplemental Figure S1-S3 for "Brain State Dynamics and Developmental Differences in Reading Comprehension"

**Supplementary Materials**

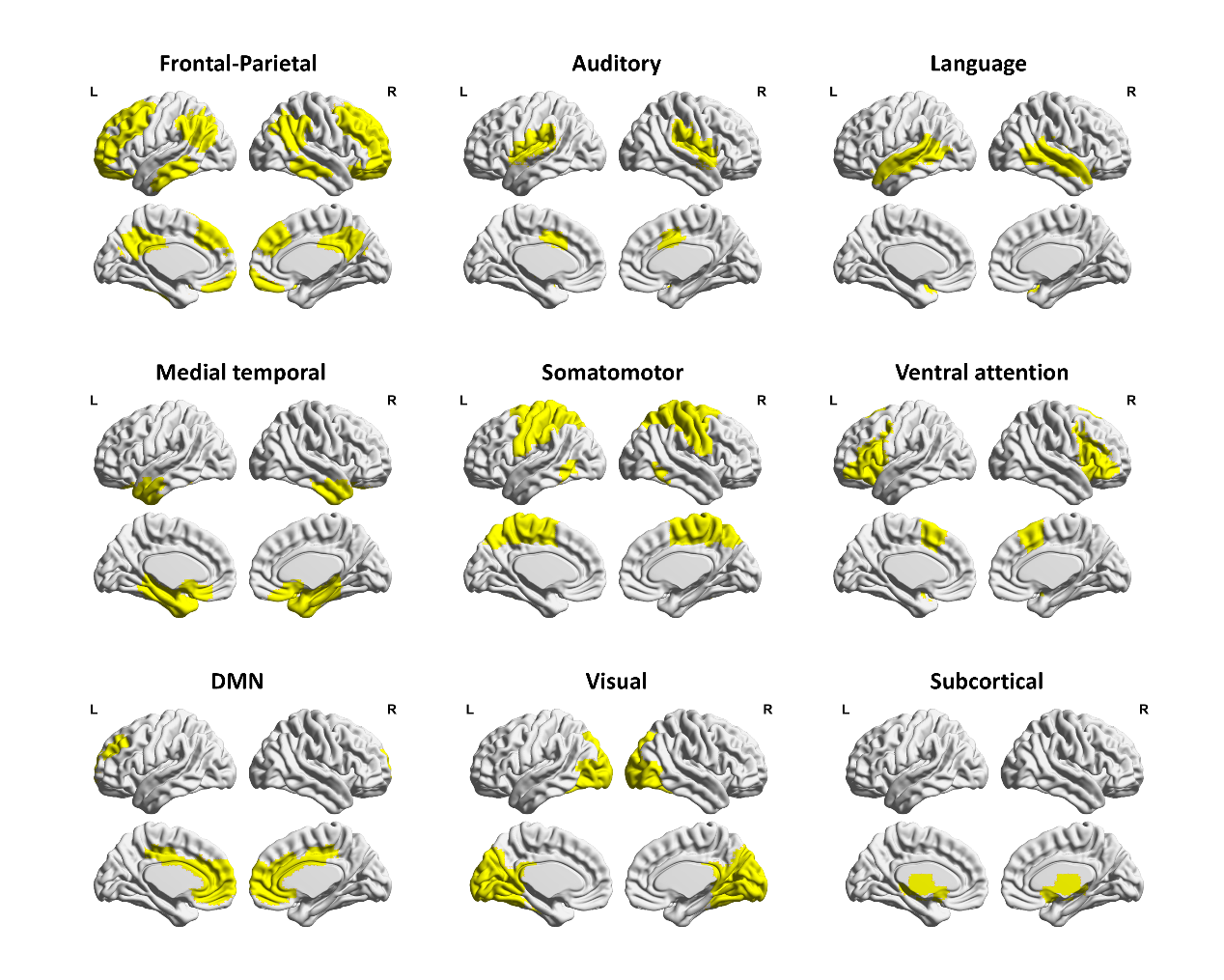

Fig. S1. The nine-network partition used in our current study.

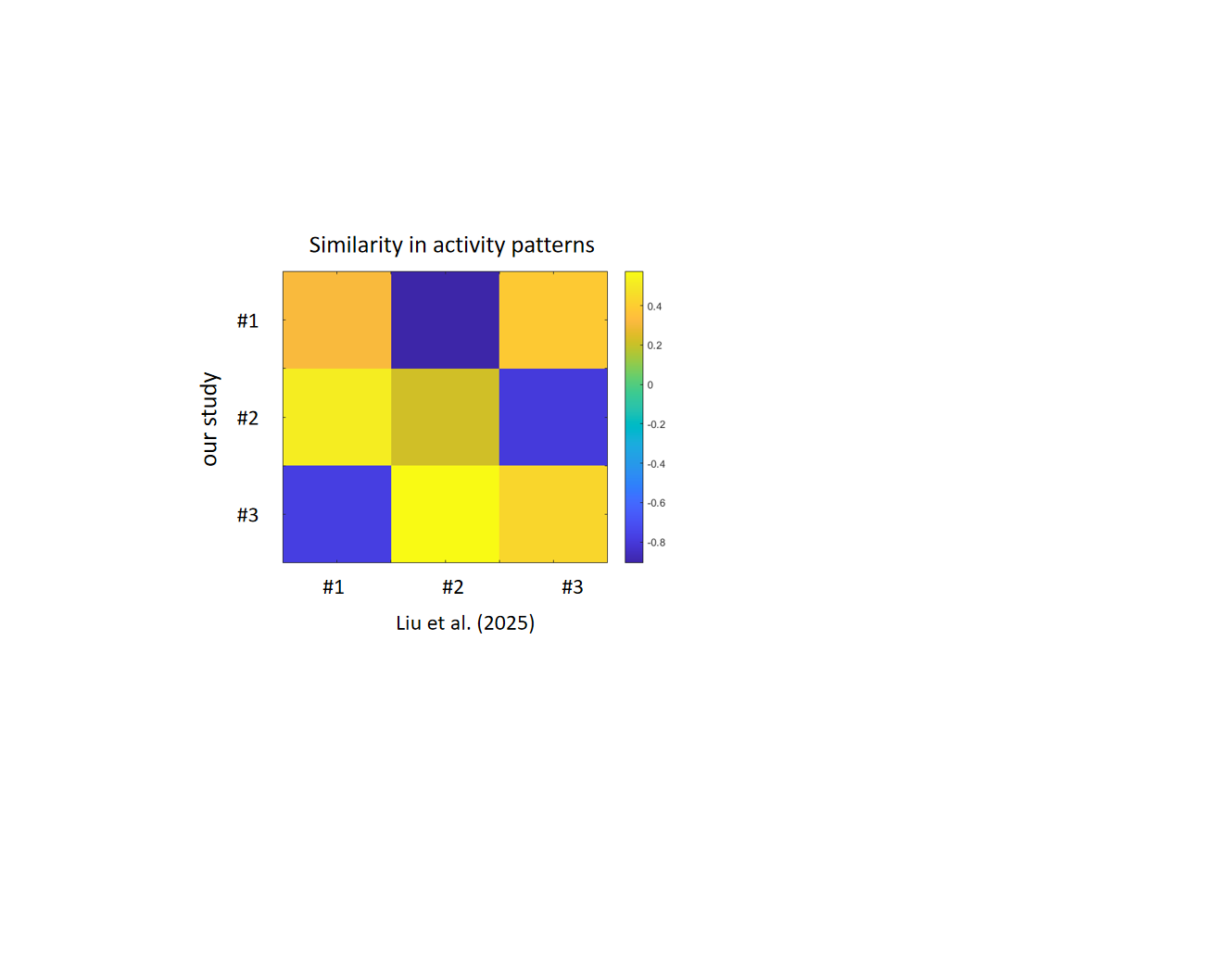

Fig. S2. The similarity of activity patterns between the three states in our study (reading comprehension task) and those reported by Liu et al. (2025) (speech comprehension task).

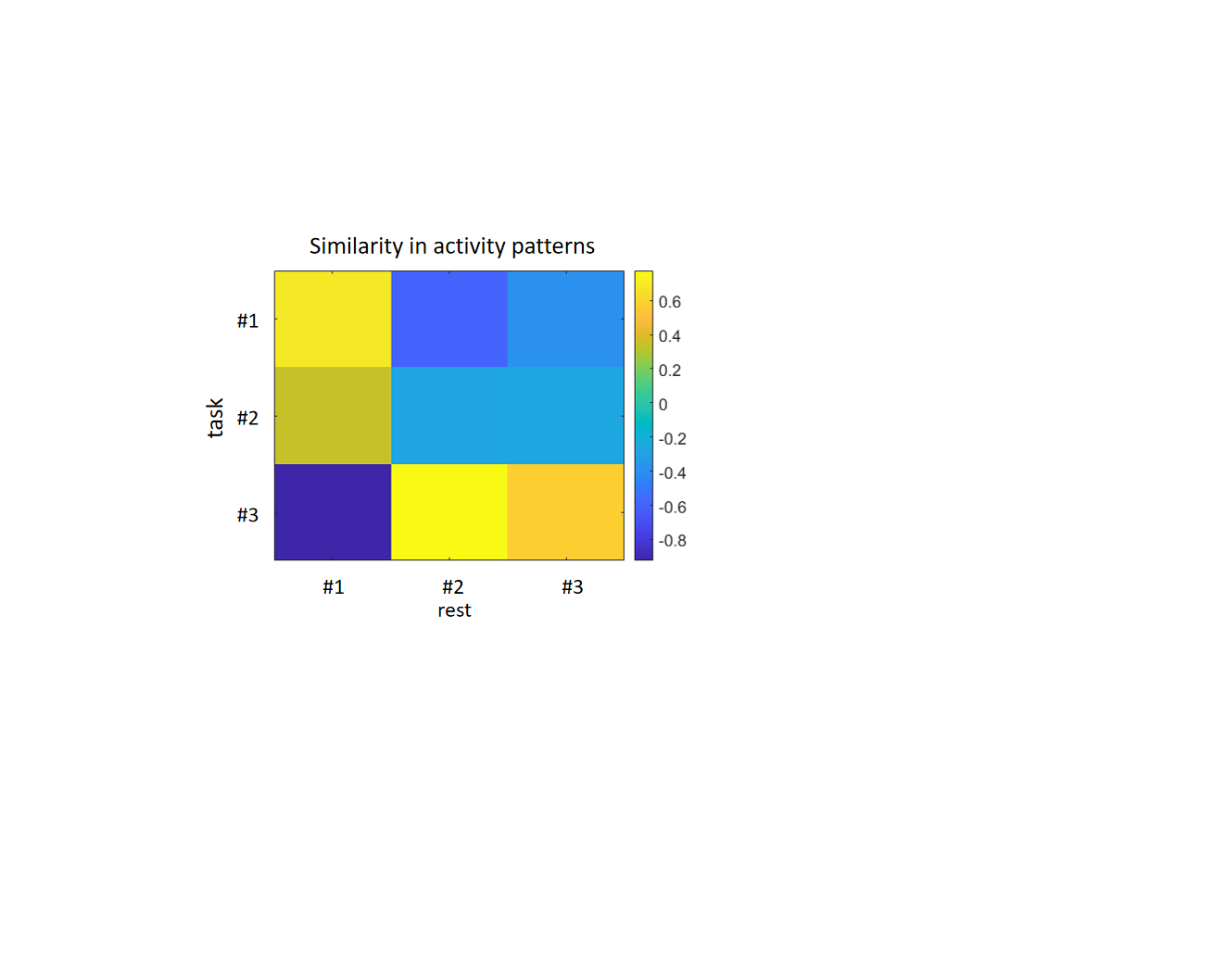

Fig. S3. The similarity in activity patterns of three states between reading comprehension task and resting state.
